# Differentiation of Wharton’s Jelly-derived mesenchymal stem cells into insulin-producing beta cells with the enhanced functional level on electrospun PRP-PVP-PCL/PCL nanofibers scaffold

**DOI:** 10.1101/2023.08.28.555005

**Authors:** Seyed Mohammad Javad Hashemi, Seyed Ehsan Enderami, Ali Barzegar, ReyhanehNassiri Mansour

**Affiliations:** Department of Basic Sciences, Sari Agricultural Sciences and Natural Resources University, Sari, Iran; Immunogenetics Research Center, Department of Medical Biotechnology, School of Advanced Technologies in Medicine, Mazandaran University of Medical Sciences, Sari, Iran; Department of Tissue Engineering, School of Advanced Technologies in Medicine, Mazandaran University of Medical Sciences, Sari, Iran

**Keywords:** insulin-producing cells, 3D culture, PRP-PVP-PCL/PCL nanofibrous scaffold, PRP, Wharton’s Jelly-derived mesenchymal stem cells, Pancreatic tissue engineering

## Abstract

Diabetes is a global problem that threatens human health. Cell therapy methods using stem cells and tissue engineering of pancreatic islets as new therapeutic approaches have increased the chances of successful diabetes treatment. In this study, to differentiate Wharton’s Jelly-derived mesenchymal stem cells (WJ-MSCs) into insulin-producing cells (IPCs) with improved maturity and function, platelet-rich plasma (PRP)-Polyvinylpyrrolidone (PVP)-Polycaprolactone (PCL)/PCL nanofiber scaffold was designed and used. WJ-MSCs-derived IPCs on PRP-PVP-PCL/PCL scaffold took round cluster morphology, which is the typical morphology of pancreatic islets. Real-time PCR, immunocytochemistry, and flow cytometry data showed a significant increase in pancreatic marker genes and insulin in WJ-MSCs-derived IPCs on the PRP-PVP-PCL/PCL scaffold compared to the two-dimensional (2D) experimental group. Also, using the ELISA assay, a significant increase in the secretion of insulin and C-peptide was measured in the WJ-MSCs-derived IPCs of the three-dimensional (3D) experimental group compared to the 2D experimental group, which indicated a significant improvement in the functional level of the WJ-MSCs-derived IPCs in the 3D group. The results showed that the PRP-PVP-PCL/PCL scaffold can provide an ideal microenvironment for the engineering of pancreatic islets and the generation of IPCs.

## Introduction

Diabetes mellitus type 1 (T1DM) is an autoimmune disorder with a decrease in the function of the pancreatic islets because of the gradual destruction of insulin-producing beta cells. This disease has the same prevalence in males and females(Warshauer et al., 2020; Perkins et al., 2021). Both genetic and environmental factors contribute to the risk of developing this polygenic disease(Saberzadeh-Ardestani et al., 2018). The result of the loss of insulin-producing cells (IPCs) is persistent hyperglycemia, which causes complications such as stroke, and heart and kidney diseases(Riddell et al., 2020). Accurate control of blood glucose at different times is not possible with exogenous insulin injections as a standard treatment for T1DM. As a result, hypoglycemia caused by continuous insulin injection is observed. Transplantation of the whole pancreas and islets of the pancreas are considered as ways to avoid daily insulin injection between these two treatment methods. Islet transplantation is ideal. In the transplantation of pancreatic islets, there are problems such as insufficient access to islet donors, the need to obtain transplanted islets from cadavers, and the low efficiency of islet transplantation(Riddell et al., 2020). The treatment method based on cell therapy through the generation of mesenchymal stem cells (MSC)-derived IPCs is considered a new treatment method and an alternative to the two previously mentioned methods(Laidmäe et al., 2022). In this method, it should obtain differentiated IPCs with a high ability to regulate the amount of insulin secretion to various concentrations of glucose(Zhang et al., 2022). To generate differentiated beta cells with morphology and function closer to the beta cells in pancreatic islets *in vivo*, it is necessary to mimic the natural microenvironment and physiology of pancreatic islets *in vitro*(Laidmäe et al., 2022). Electrospun nanofiber scaffolds (ENFs) are one tool used for this imitation. In the design of these scaffolds, the extracellular matrix (ECM) and microenvironment of cells play an important role in cell-cell and cell-ECM interactions and these interactions are necessary for the formation of island clusters(Sánchez-Cardona et al., 2021; Laidmäe et al., 2022; Zhang et al., 2022). ENFs have a structure similar to ECM(Laidmäe et al., 2022; Mirzaei et al., 2019). In the fabrication of these scaffolds, synthetic materials, natural materials, or both are used(Bhattarai et al., 2018). Polycaprolactone (PCL) is a biodegradable and biocompatible chemical polymer that is used in the fabrication of ENFs(Himmler et al., 2022). Polyvinylpyrrolidone (PVP) is a water-soluble(Haaf et al., 1985). And non-toxic, and non-ionic chemical polymer(Lu et al., 2012; Jadhav et al., 2013). It is widely used in the medical industry and the fabrication of various types of ENFs. Nanofiber scaffolds made of PVP can carry a variety of biological materials(Sun et al., 2010; Liu et al., 2021). Wharton’s Jelly mesenchymal stem cells (WJ-MSCs) in the umbilical cord are good candidates for cell therapy(Maldonado et al., 2017; Musiał-Wysocka et al., 2019). These cells can be obtained easily and in large quantities by a non-invasive method from the umbilical cord, which is discarded after delivery. WJ-MSCs are derived from a proteoglycan matrix called Wharton’s Jelly in the umbilical cord tissue(Maldonado et al., 2017; Wang et al., 2019). They have the potency of proliferation and self-renewal higher than other MSCs derived from other tissues. They also have the potency of immunosuppression, low immunogenicity, and immunomodulatory properties(Maldonado et al., 2017; Musiał-Wysocka et al., 2019; Rizal et al., 2021). The very low expression of human leukocyte genes (HLA) class L, also, lack of the expression of class L genes (HLA-DR) genes cause a decrease in allo reactivity in these cells. They also lack the expression of antigens necessary for the activate B and T cells, such as CD80 and CD86(Choi et al., 2020). These cells do not have tumorigenic activity. Also, they have kept some characteristics of embryonic stem cells, such as Sox2, Oct4, c-Kit, and Nanog gene expression(Nekanti et al., 2010; Yang et al., 2021). These features make this type of cell suitable for allogeneic and autogenic transplantation(Rizal et al., 2021). The concentration of platelets in the platelet-rich plasma (PRP) obtained from autologous blood plasma is five times higher than the normal concentration(Marx, 2004). The platelets contain many and diverse growth factors which are involved in cellular processes, such as angiogenesis, differentiation, proliferation, and migration(Cakir et al., 2019). For example, platelet-derived growth factor (PDGF), fibroblast growth factors (FGF), basic fibroblast growth factor (bFGF), insulin-like growth factor 1 (IGF-1), and transforming growth factor beta (TGF-β), lead to increased cell proliferation, ECM synthesis is regulated by IGF, PDGF, and TGF-β and angiogenesis increases with hepatocyte growth factor (HGF), FGF-2, and vascular endothelial growth factors (VEGF)(Amable et al., 2013; Pötter et al., 2021; Abazari et al., 2019). In this study, we innovatively converted PRP into fiber form, which led to its stepwise release from the PRP-PVP-PCL/PCL scaffold. Then we used this *in vitro* 3D culture system to obtain WJ-MSCs-derived IPCs with round morphological characteristics and cluster cell accumulations like beta cells in pancreatic islets *in vivo*, as well as having improved functional characteristics of insulin production and high ability to regulate the amount of insulin secretion in response to different glucose concentrations.

## Materials and methods

### _ Preparation of PCL and PRP-PVP PCL solutions

PCL (15% w/v), (80,000, sigma) solution was prepared using chloroform (Dr. mojallali) solvent in a final volume of 5 ml for 5 hours at a temperature of 25^°^C on a heater stirrer (800 rpm). PRP (8% w/v) was dissolved in 96% methanol solvent, also PVP (20% w/v, PVP K90 sigma), was dissolved in dichloromethane (DCM), (Dr. mojallali) solvent, (both with an overnight at 30°C and 800 rpm), and PRP solution was added drop by drop to the PVA solution. Methanol and dichloromethane with a ratio of 4:1 were used to prepare the PRP-PVP solution. In the following, PCL (5% w/v) solution was also prepared with methanol solvent, then PRP-PVP (8%: 20%) solution was added drop by drop to PCL (5%) solution in a ratio of 1:3, for 5 hours and 1000 rpm.

### _ Electrospinning process of nanofiber scaffolds

The final volume of the PRP-PVP-PCL (8%:20%:5%) solution was 5 ml. PCL (15%) solution with a final volume of 5 ml was entered into a 16 G gauge glass syringe and PRP-PVP-PCL (8%:20%:5%) solution with a final volume of 5 ml was entered an 18 G glass syringe. Then the fabrication of the nanofiber scaffold was done by the co-electrospinning method (FNM CO. Ltd). Making PRP-PVP-PCL/PCL scaffold (8%:20%:5%/15%) was started by injecting two syringes at the same time. The injection speed in both syringes was 0.2 ml/hour, the distance between the syringes and the collector was 15 cm, the rotation speed of the rotator was 150 rpm, and the voltage of the electrospinning machine was set at 15 volts.

### _ Preparation method of PRP

Blood samples were collected from 4 women aged 30 to 40 to prepare PRP. The amount of 10 ml of blood from each donor became centrifuged at 2000 rpm, for 15 min. Plasma and buffy coat layers had been amassed and half of the plasma turned into removed after the next centrifugation (4000 rpm, 18 min). Platelets were activated with CaCl2 (10%) and after overnight storage at 4 °C were filtered and geared up to use(Pakfar et al., 2017). Next, the process of lyophilization of the PRP solution was conducted for ease of storage and fiberization.

### _ Sterilization of fabricated nanofiber scaffolds

Initially, discs with a diameter of 1.5 cm were prepared from the synthesized nanofiber scaffold (PRP-PVP-PCL/PCL) and placed on a 24-well cell culture plate. Next, filtered 70% ethanol was added to the discs inside each well, and after 24 hours of immersion in 70% ethanol, washing was done three times with a phosphate buffer solution (PBS). Finally, by adding a complete cell culture medium to each well, incubation was done for 24 hours to ensure the absence of microbial contamination.

### _ Scaffolding characterization tests

### _ Degradation Test

Samples of PVP-PCL and PRP-PVP-PCL nanofiber scaffolds with specific dimensions (10 × 10 mm) were placed in vials containing 4 ml of PBS (pH = 7.40 at 37^◦^C) for 3 days. Three replicates were placed for each scaffold sample. The PBS solution was replaced three times a day and at certain times. The parameters of water absorption rate and mass loss rate of nanofiber scaffolds were investigated for the scaffold degradation test. The weight of the samples was measured in 1,2 and 3 days and the percentage of degradation of the scaffolds was measured. Wd is the final weight (mg) of nanofiber scaffold samples. Wt is the weight of the dry samples of the nanofiber scaffold

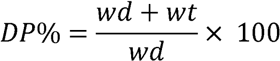

### _ PRP release assay from nanofiber scaffold

Fragments of PRP-PVP-PCL/PCL scaffolds with specified dimensions have been positioned inside the PBS solution for 14 days. To measure total protein, at certain times every day, 0.5 ml of PBS solution was removed to measure the total protein in the solution, and 0.5 ml of fresh PBS was replaced again. BCA assay kit (Pierce, Rockford, IL, USA) was used to measure the amount of protein. The assay was performed according to the kit instructions.

### _ Infrared spectroscopy analysis

Fourier transform attenuated total infrared reflection (FTIR-ATR) became used for the structural evaluation of the scaffolds, the use of a spectrum one spectrometer (Perkin-Elmer, Shelton, CT, USA). All spectra were taken within the spectral variety of 4000–500 cm^-1^ with a resolution of 4.0 cm^-1^ and 32 scans. The spectra have been normalized with the use of the spectrometer software program (SPECTRUM, v.10.5.3, Perkin-Elmer Inc., 2016, Shelton, CT, USA) with an ordinate restrict of up to at least 1.0 absorbance.

### _ Contact angle analysis

A drop of water was placed on each of the PCL, PRP-PVP, and PRP-PVP-PCL scaffolds and after 10 seconds, the contact angle for each scaffold sample was analyzed using the contact angle protractor (GLOKRUSS).

### _ Survival evaluation

To measure the livability of the cells on the fabricated nanofiber scaffolds and confirm the non-cytotoxicity of the scaffolds, samples of PVP-PCL, PVP-PRP, and PRP-PVP-PCL scaffolds with the number of three replicates for each sample were placed in the cell culture plate which contained complete cell culture medium and incubated (37^°^C, 95% humidity, and 5% CO2). After 72 hours, the medium on the scaffolds, which contained the components of each type of scaffold, was removed to assay the cell viability and non-cytotoxicity of the fabricated scaffolds, and it was transferred to the wells of a 12-well cell culture plate. WJ-MSCs were added in the number of 4 × 10^3^ cells/cm^2^ to each of the culture mediums treated with scaffolds and incubated. Three replicates were considered as an untreated control group. After 1, 3, 5, and 7 days of incubation, the medium on the cells attached to the bottom of the wells was removed and the MTT test was performed.

### _ Isolation of Wharton’s Jelly-derived mesenchymal stem cells by explant method

The study was approved by the Ethics Committee of the Mazandaran University of Medical Sciences (Code: IR.MAZUMS.4. REC.1402.17642). Initially, a piece of the umbilical cord after cesarean section is placed in physiological serum after obtaining legal consent from a healthy volunteer mother and is transferred from the imam hospital (Sari, Iran) to the laboratory. Then it is disinfected under a class L laminar hood with 70% ethanol for 30 seconds. Then the umbilical cord sample was placed in Hank’s balanced salt solution (HBSS) and cut into 2 cm sections. Then, the umbilical cord epithelium vessels were gently separated. Pieces of Wharton’s Jelly were gently removed and transferred into the HBSS dish. Wharton’s Jelly tissue was minced with a sterilized scalpel. Minced pieces were cultured in a T25 flask which contained DMEM/F12 (GIBCO, UK) (Dulbecco’s Modified Eagle’s Medium) and FBS (fetal bovine serum) (SIGMA-Aldrich) 15% and 1% antibiotic (penicillin/streptomycin). The flask was transferred to an incubator (37^°^C, 95%, 5%), and after 3 days, the culture medium of the flask was changed(Mirtaghi et al., 2022).

### _ Characterization of mesenchymal stem cells by flow cytometry

The cultured WJ-MSCs in passage number 3 were trypsinized and, after washing, they were exposed to CD166, CD73, CD90, CD105, CD45, and CD34 antibodies and incubated for half an hour at 4^°^C. Then, the expression or lack of expression of these CD markers was analyzed using flow cytometry.

### _ Cell seeding on scaffolds and experimental design

Initially, discs with a specific diameter were prepared from the PRP-PVP-PCL/PCL scaffold and placed on a 24-well cell culture plate. And after washing three times with PBS, they were incubated with DMEM/F-12 (Dulbecco’s Modified Eagle Medium/Nutrient Mixture F-12) (GIBCO, UK) for 24 hours. 2D and 3D PRP-PVP-PCL/PCL control groups, also 2D and 3D PRP-PVP-PCL/PCL experimental groups were considered to continue the differentiation process. Then about, 50,000 WJ-MSCs were added to each well in all groups with the full medium which contained DMEM/F-12 (GIBCO, UK) and FBS (SIGMA-Aldrich) 15% and 1% antibiotic (penicillin/streptomycin) for 3 days, and when they reached a density of 70-80%, differentiation medium was added in two steps to the experimental groups and the differentiation process was done in 20 days. In the first stage of the cell differentiation process, the differentiation culture medium L which contained DMEM/F-12 medium supplemented with 2% B27 (Invitrogen), 10 ng/ml epidermal growth factor (EGF; Sigma), 10 ng/ml basic fibroblast growth factor (b FGF; Sigma), 10 mmol/L nicotinamide (Sigma), 2% FBS was added to the 2D and 3D experimental groups and were incubated (temperature of 37^°^C, 95% humidity, and 5% CO2) for 10 days. In the second step of the differentiation process, differentiation medium L which contained DMEM/F-12 medium supplemented with 10 ng/ml EGF (Sigma), 50 ng/ml activin A (Sigma), 10 mmol/L nicotinamide (Sigma), 2% B27, 10 ng/ml exendin-4 FBS was added to both experimental groups and incubation was done for another 10 days. During the 20 days of the differentiation process, the cell culture medium in the both experimental and control groups was changed with a fresh medium every two days. All control groups were without differentiation culture medium and contained full culture medium.

### _ Scanning electron microscopy (SEM)

Cellular PRP-PVP-PCL/PCL nanofiber scaffolds were stabilized in 2.5% glutaraldehyde (Merck, Darmstadt, Germany) for 45 min. Then the stabilized samples were washed twice with PBS for 5 min. Then dehydration was done for 10 minutes in each of the 50, 60, 70, 80, 90, and 100% v/v, ethanol series. With Araldite TM glue, the dried scaffolds were mounted on the holder and coated in a sputter-coater (Hitachi, Tokyo, Japan) with palladium gold. Then, imaging of the scaffolds was done with a scanning electron microscope (SEM; Hitachi S-4500, Japan) at an accelerating voltage of 5 kV.

### _ Real-time PCR analysis

Initially, total RNA was extracted from about 1 × 10^6^ cells from all groups using RNeasy Mini Kit (Qiagen, Valencia, CA), then using the random hexamer primer and M-MuLV reverse transcriptase kit (Fermentas, Helsingborg, Sweden), cDNA synthesis was performed according to the instructions of the kit. The real-time PCR reaction was performed using an ABI Step One system and using SYBR premix ExTaq, (Takara Bio, Shiga, Japan). Data normalization was done with beta-2-microglobulin (β2M) as a reference gene and by using the relative quantification method (2^-ΔΔCt^) data were analyzed. The primers are shown in table 1.

### _ Immunocytochemistry assay

Initially, the cells of all groups were washed twice with PBS. Then they were stabilized with 4% cold paraformaldehyde (Sigma Aldrich) for 20 minutes at 4^°^C and 5 minutes at room temperature (RT). Then they were permeabilized with 0.2% Triton X-100 (Sigma Aldrich). Next, the blocking buffer containing goat serum was used for 45 minutes at RT. After removing the supernatant, the cells were exposed to the primary antibody (anti-insulin, 1:200; #ab181547) overnight at 4°C. Next, the cells were washed three times with 0.1% Tween at 5-minute intervals. Cells were blocked with 1% bovine serum albumin (BSA) and incubated for 30 minutes at RT. The supernatant solution was removed and the cells were treated with a secondary antibody (the FITC-conjugated anti-mouse IgG (Sigma)) for one hour at 37°C in the darkness. After washing three times with 0.1% Tween for 5 minutes, nuclear staining was done with 0.1 μg/ml DAPI (4′,6-diamidino-2-phenylindole) (Sigma-Aldrich) for 1 minute, followed by washing with PBS(Enderami et al., 2017). For secondary control, the IPCs were incubated with secondary antibody, without primary staining. Images were taken with a phase contrast fluorescent microscope (Nikon, Tokyo, Japan). Immunocytochemistry (ICC) images were quantified by ImageJ software (Version 1.45 s, NIH, USA).

### _Flow cytometry assay of insulin

At first, in all groups, cells were washed twice with PBS. After being suspended with 4% paraformaldehyde fixative, they were incubated for 20 minutes at RT. Cell permeability was done using 0.4% Triton X-100, after washing the cells twice with PBS. Then they were diluted with PBS. It was done for 1 hour. Then, blocking was done using normal goat serum. After adding 10 µl FITC-conjugated anti-insulin antibody (R&D system), incubation was done at RT and away from light for one hour. After washing twice with Tween buffer, they were suspended in 300 µl of PBS, and evaluation was done using flow cytometry (Attune Acoustic Focusing Cytometer) and FlowJo software.

### _Insulin and C-peptide release in response to glucose stimulation

The cells of all the control and 2D and 3D experimental groups were incubated in glucose-free Krebs-Ringer bicarbonate (KRB) buffer containing 5 mM KCl, 25 mM NaHCO3, 2.5 mM CaCl2, 120 mM NaCl, 1.1 mM MgCl2 and 0.1% BSA (Sigma Aldrich) for 2 hours. Then they were incubated in different concentrations of glucose for 30 minutes, and then the medium was collected and the amount of insulin and C-peptide secretion was measured. It was measured with the ultrasensitive ELISA kit (Mecodia, Uppsala, Sweden) based on the kit protocol.

### _Statistical Analysis

After at least three independent repetitions of each test, the results were reported as a mean-± standard deviation (SD). The analysis of the obtained results was done by one-way analysis of variance (ANOVA). Comparison between 2D and 3D groups was done with Bonferroni’s Posthoc test by GraphPad Prism 6 software (Graphpad Software Inc., La Jolla, CA). A value of p < 0.05 was considered statistically significant.

## Results

### _Characterization of isolated WJ-MSCs

To confirm the mesenchymal nature of cells isolated from Wharton’s Jelly tissue, after the third passage, the expression of surface markers of MSCs was evaluated by flow cytometry. Flow cytometry results showed high expression of CD105 (99.4), CD73 (93.8), CD90 (99.0), and CD166 (81.2) as specific WJ-MSCs markers. Also, the very low expression of CD34 (0.478), and CD45 (0.896) markers, which are hematopoietic specific, was confirmed (Fig. 1).

**Fig. 1.**
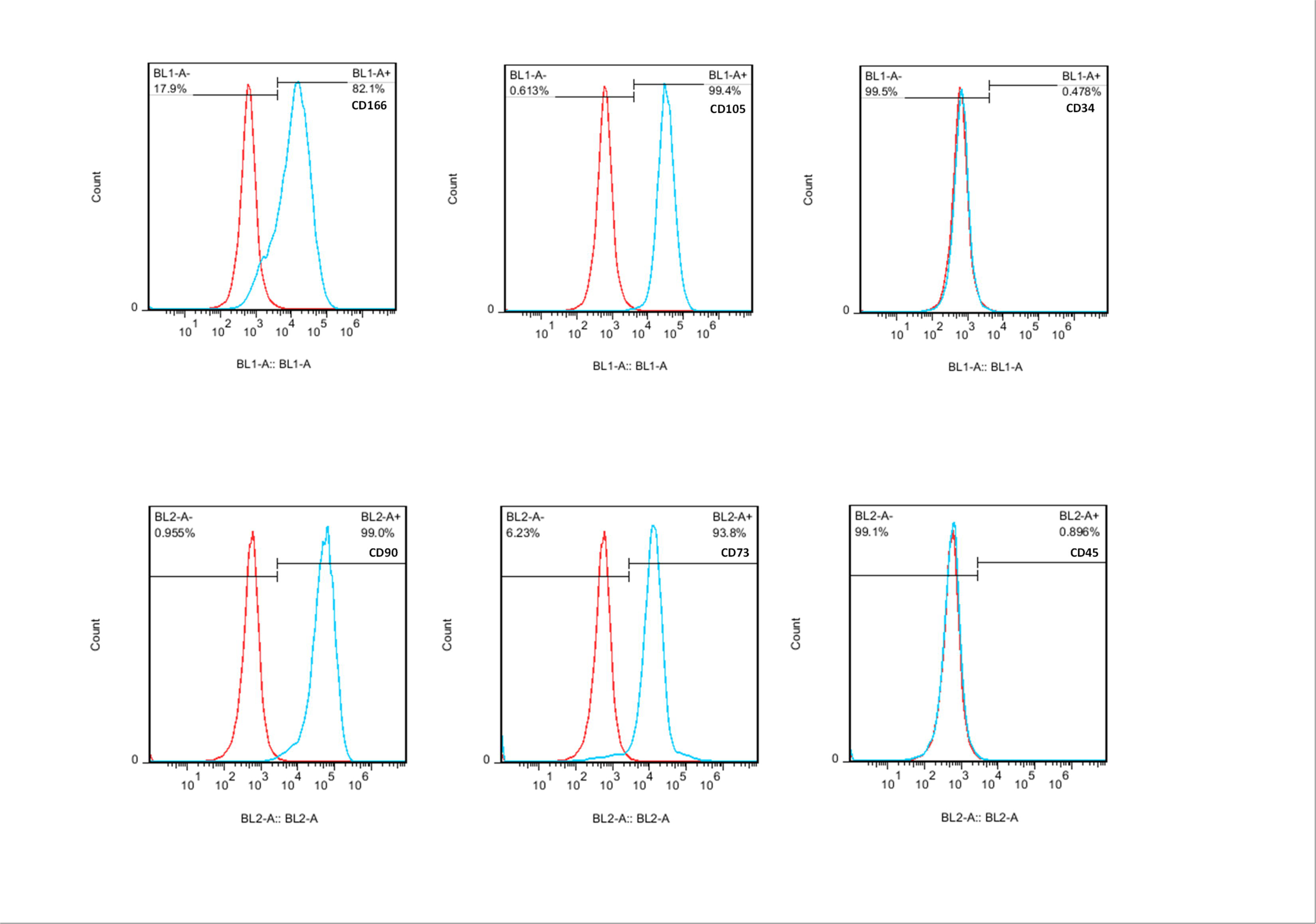
Flow cytometry results for isolated WJ-MSCs. The cells were positive for CD105, CD90, CD73, and CD166 markers and negative for CD34 and CD45 markers.

### _Evaluation of cell morphology changes in 2D and 3D cell cultures

After completing the two-step differentiation protocol, the images of the cells of the 2D culture groups were recorded with an inverted microscope (Fig. 2A, B), and the images of the cells of the 3D culture groups were recorded with the SEM (Fig. 2C, D). The cells of the 2D control group (Fig. 2A) that were cultured in a non-differentiated culture medium were unchanged and they were spindle-shaped and fibroblast-like structures. Also, they did not form island-like structures. But the WJ-MSCs cultured in the differentiation medium after the differentiation period (Fig. 2B), had a spherical shape like IPCs, and by gathering together, they formed islet structures similar to pancreatic islets of Langerhans. In 3D groups (Fig. 2C, D), cells settled in the pores of the nanofiber scaffolds, in the control group (Fig. 2C), they have a flat and elongated morphology. Also, in the 3D experimental group (Fig. 2D), after the differentiation period, they obtained a round morphology similar to IPCs and gathered as spherical clusters similar to pancreatic islet clusters.

**Fig. 2.**
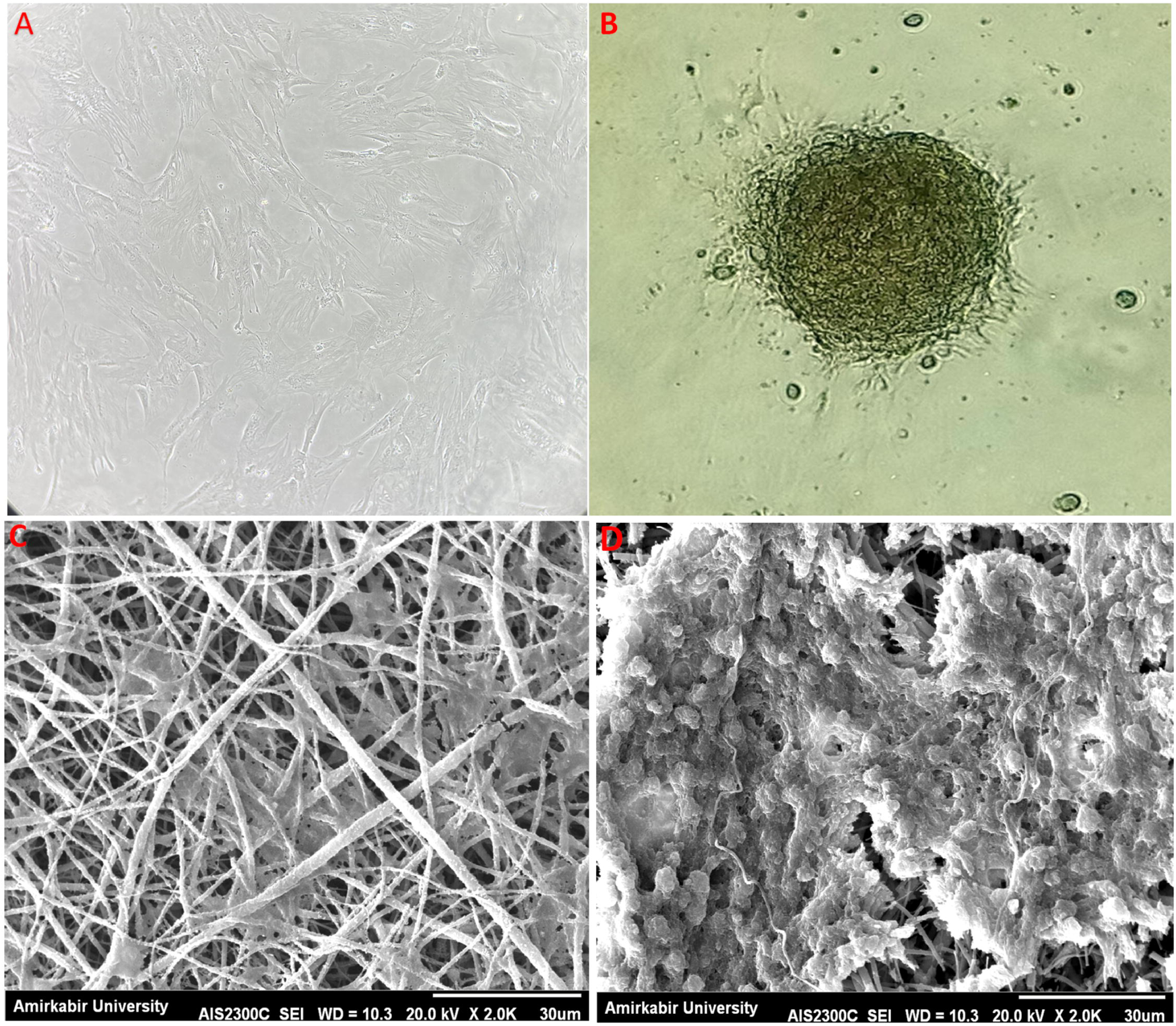
Morphological changes of cells in 2 D and 3D culture. WJ-MSCs in undifferentiated culture medium (2D control group), WJ-MSCs were spindle-shaped and similar to fibroblast cells (A) WJ-MSCs-derived IPCs in differentiation culture medium, the spherical WJ-MSCs-derived IPCs formed cellular aggregates similar to pancreatic islets. Scale bars are 100 µm (B). WJ-MSCs seeded on the scaffold (PRP-PVP-PCL/PCL) in an undifferentiated culture medium (3D control group), The cells have a spindle-shaped morphology(C). WJ-MSCs -seeded scaffold (PRP-PVP-PCL/ PCL scaffolds) in IPC differentiation medium. WJ-MSCs-derived IPCs formed dense spherical and round clusters. Scale bars are 30 µm (D).

### _Scaffold characterization

After the fabrication of nanofiber scaffolds, the degradation percentage of scaffolds was tested, and the results showed that, after 72 hours, PRP-PVP-PCL/PCL, PVP-PCL, and PCL scaffolds lost 83%, 47%, and 2.5% of their mass, respectively (Fig. 3A). The degraded part of the mass of the scaffolds was related to their structural hydrophilic parts (PVP and PRP). With the increase of hydrophilicity in the composition of the scaffolds, the mass degradation percentage increased. The evaluation of the cumulative release profile of PRP from the PRP-PVP-PCL/PCL nanofiber scaffold showed the burst release of PRP during the first 48 hours. Then, the release of PRP from the nanofiber scaffold continued with a slow-increasing pattern until the end of the second week (Fig. 3B). Contact angles for PCL, PVP-PCL, and PRP-PVP-PCL scaffolds were reported as 92^0^, 61^0^, and 33^0^ degrees, respectively (Fig. 3C). PCL showed the highest contact angle because of its hydrophobic surface, after adding PVP to the scaffold composition and then adding PRP, the hydrophilicity property on the surface of the scaffold increased, further increase in the hydrophilicity property on the surface caused a further decrease in the contact angle. In the fabrication of nanofiber scaffolds, the preservation of the chemical nature of the components after combining them was confirmed by FTIR analysis (Fig. 3D). PCL and PVP used in our research had 2945 cm^-1^ (-CH2-),1723 cm^-1^(–C = O) and 1286 cm^-1^(–C-H-),1373 cm^-1^(-CH2-), 2921 cm^-1^(-CH2-) peaks, respectively, which confirmed their chemical nature. PVP-PCL scaffold made in our research had peaks of 1723 cm^-1^ (–C = O) and 2919 cm^-1^ (-CH2-), which confirmed the preservation of the chemical nature of PVP and PCL after mixing. After that, the PRP-PVP-PCL scaffold was fabricated by adding PRP. Peaks 2945 cm^-1^ (-CH2-), 1723 cm^-1^ (–C = O), and 1647 cm^-1^ (–C = O) confirmed the preservation of the chemical nature of PCL and PVP, also peaks 1532 cm^-1^and 1257 cm^-1^, which correspond to Amid L and Amid L respectively, pointed to the proteins in PRP. Also, we had a peak in 3327 cm^-1^ related to the (N-H) group in the proteins of PRP.

**Fig. 3.**
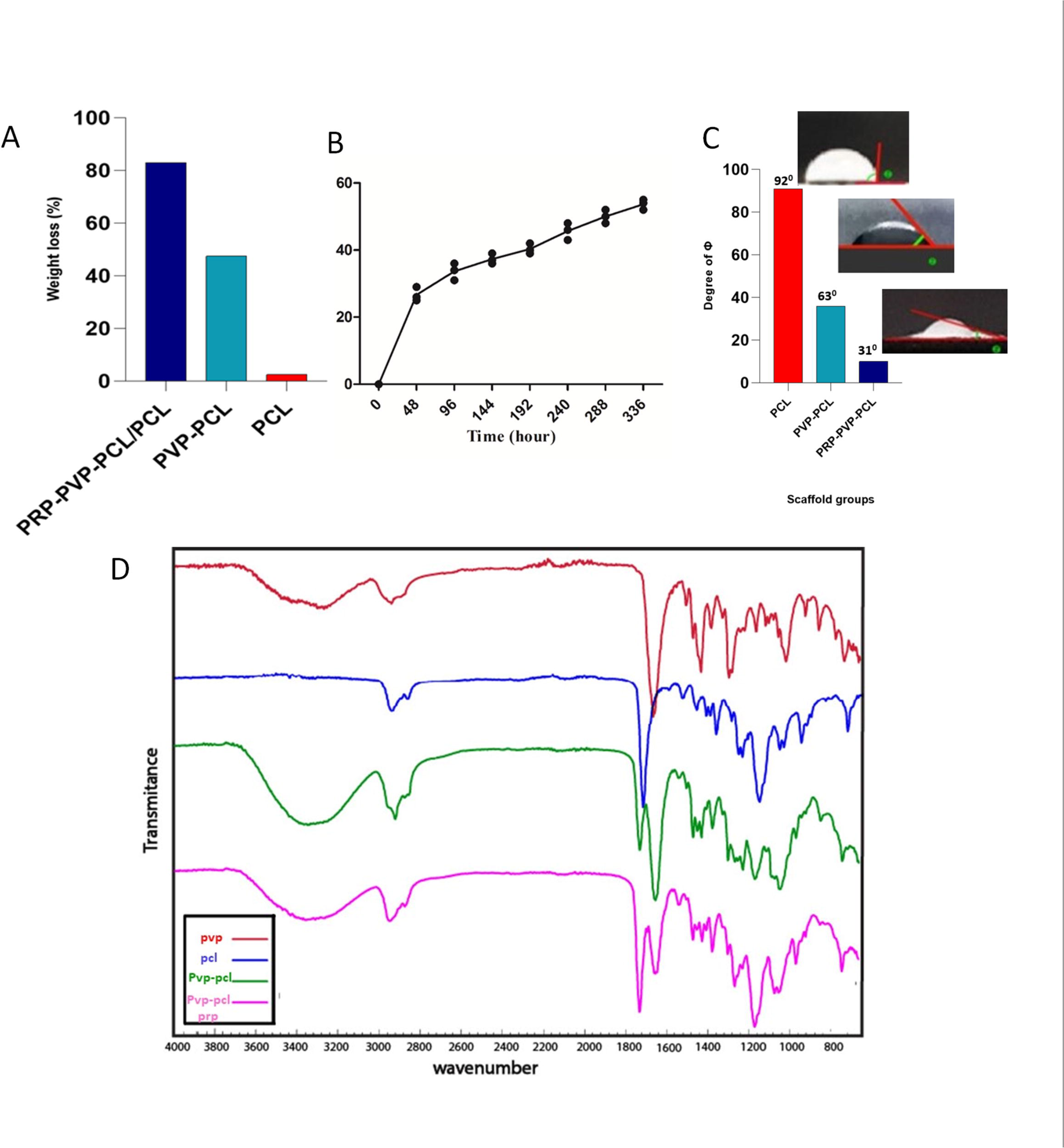
Characteristics of fabricated nanofiber scaffolds. Degradation percentage for PRP-PVP-PCL/PCL, PVP-PCL, and PCL scaffolds after 72 hours. PRP-PVP-PCL/PCL, PVP-PRP, and PCL scaffolds lost 83%, 47%, and 2.5% of their mass, respectively (A). The cumulative release (%) of PRP from PRP-PVP-PCL/PCL scaffold after 14 days. The burst in PRP release occurred after 48 hours (B). Contact angle test for PCL, PVP-PCL, and PRP-PVP-PCL scaffolds. PRP-PVP-PCL, PVP-PCL, and PCL respectively showed more reduction in contact angle (C). FTIR spectrum of PVP, PCL, PVP-PCL, PVP-PRP-PCL scaffolds (D).

### _Biocompatibility assay of nanofiber scaffolds

3D scaffolds should be biocompatible and not cause any cytotoxicity after cell seeding. After seeding the WJ-MSCs on the scaffolds in 3D groups, as well as their cultures in the 2D group, the biocompatibility of the scaffolds was evaluated by the MTT ([3-(4,5-Dimethylthiazol-2-yl)-2,5-Diphenyltetrazolium Bromide]) test (Fig. 4). Based on the results of the MTT test, the survival rate and cell proliferation in the 3D groups were higher than in the 2D group, but it was not significant. On the other hand, PRP-PVP-PCL/PCL scaffold showed non-significantly higher biocompatibility than the PVP-PCL scaffold. These results showed that the combination of PVP and PCL in the PVP-PCL scaffold does not cause any cytotoxicity. It was further confirmed that the addition of PRP to PVP and PCL for the fabrication of the PRP-PVP-PCL/PCL scaffold did not cause any cytotoxicity.

**Fig. 4.**
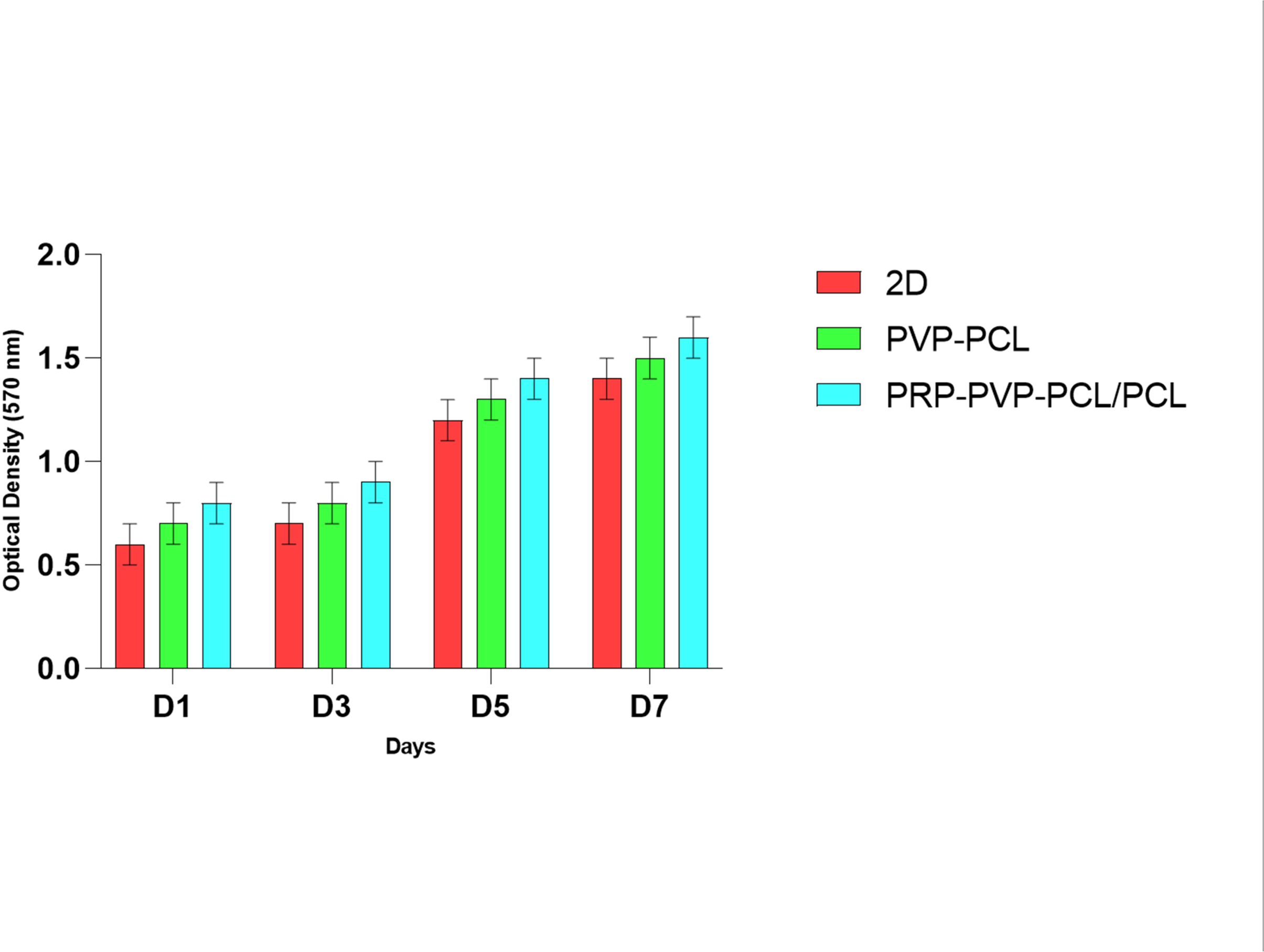
MTT assay of WJ-MSCs on PVP-PCL, PRP-PVP-PCL scaffolds, and 2D culture (control) during 1, 3, 5, and 7 days. In the 3D groups, a non-significant increase in cell survival was observed compared to the 2D control group.

### _Real-time PCR assessment

At the end of the 20th day of the differentiation period, total RNA was extracted from all studied cell groups. Then, the expression of pancreatic marker genes was compared between the 3D and 2D experimental groups with the real-time PCR, to assay the effectiveness of the PRP-PVP-PCL/PCL scaffold in promoting the differentiation protocol in this study. In (Fig. 5), the relative gene expression of the insulin, glucagon (GCG), glucose transporter 2 (GLUT2), and pancreatic and duodenal homeobox 1 (Pdx1) genes at the mRNA level are shown. In undifferentiated WJ-MSCs in the control group, the mentioned genes had a low expression level. But they had a high level of expression in 2D and 3D experimental groups. The expressions of these genes in the WJ-MSCs-derived IPCs in the 3D group (PRP-PVP-PCL/PCL), were significantly higher than the WJ-MSCs-derived IPCs in the 2D group (***P<0.001).

**Fig. 5.**
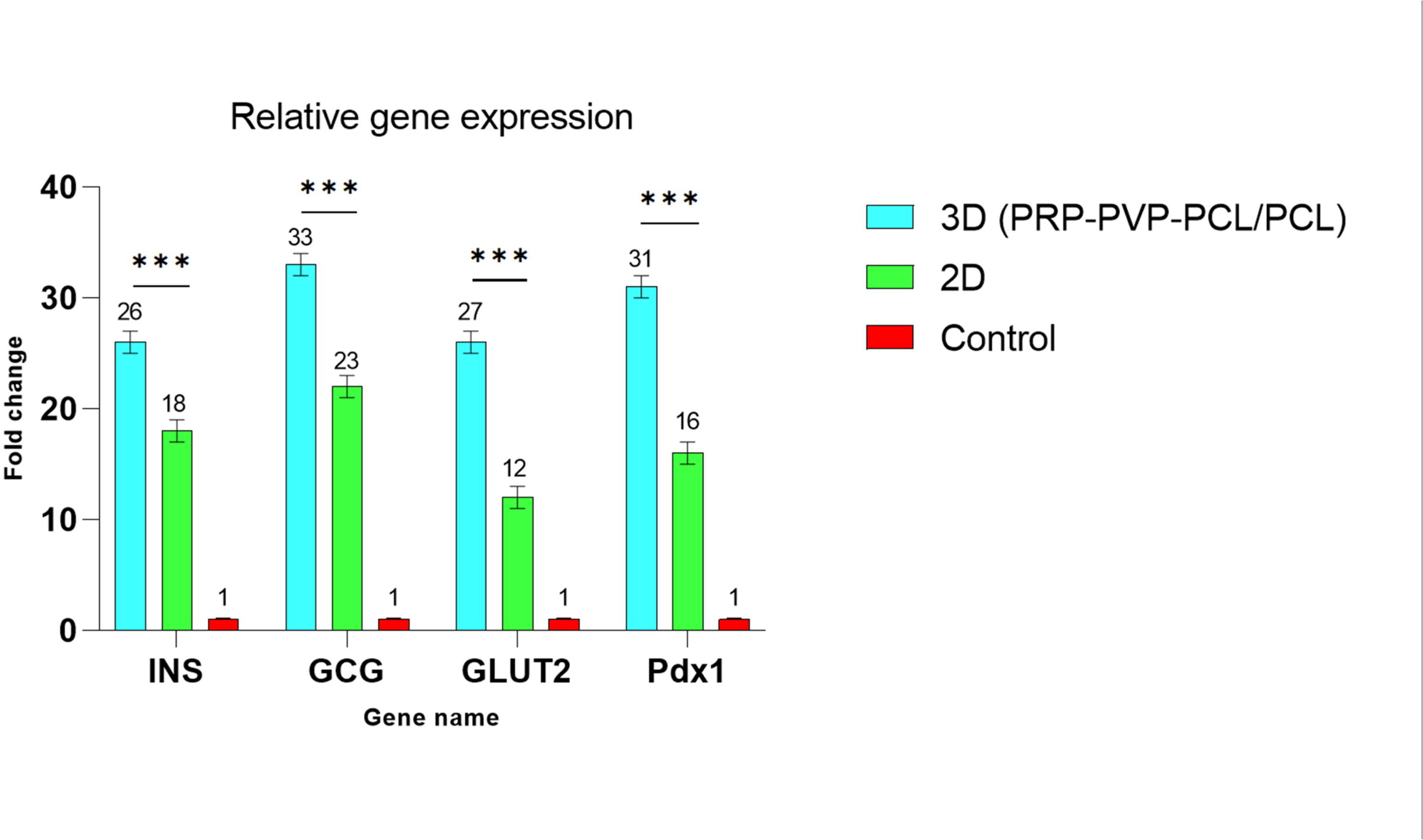
Comparison of the relative expression of pancreas-specific genes in WJ-MSCs-derived IPCs after the end of the cell differentiation period, between 2D and 3D groups. Relative gene expressions were normalized to the human β2M as a reference gene. The value is shown in each graph as mean ± SD. ***P<0 .001

### _Immunocytochemistry and flow cytometry assay

To confirm the differentiation of WJ-MSCs into IPCs, qualitative analysis of insulin gene expression at the protein level was evaluated with ICC (Fig. 6). The results of ICC confirmed the expression of insulin as a specific marker of IPCs at the protein level. WJ-MSCs-derived IPCs in both 2D (Fig. 6A1) and 3D (Fig. 6A2) experimental groups have expressed insulin protein. Also the 2D secondary controls were analyzed only by incubating IPCs with secondary antibody (Fig. 6A3) and staining of nucleus (blue) was performed by DAPI in 2D, 3D groups (Fig. 6B1, B2) and along with 2D secondary control assay (Fig. 6B3). Quantification of ICC images was done by ImageJ software (Fig. 6C) and there was a significant difference between the 2D and 3D groups (*p < .05) . After the qualitative evaluation of insulin expression at the protein level in WJ-MSCs-derived IPCs in both 2D and 3D experimental groups with the ICC test, quantitative analysis of insulin expression at the protein level in WJ-MSCs-derived IPCs was performed by flow cytometry assay (Fig. 6D1, D2). The results of flow cytometry showed that 51% of the cells in the 3D experimental (Fig. 6D1) group that was seeded on the PRP-PVP-PCL/PCL scaffold were WJ-MSCs-derived IPCs, and in the 2D experimental group (Fig. 6D2), 43% of the cell population were WJ-MSCs-derived IPCs. As a result, the 3D scaffold has increased the population of WJ-MSCs-derived IPCs in 3D culture compared to 2D.

**Fig. 6.**
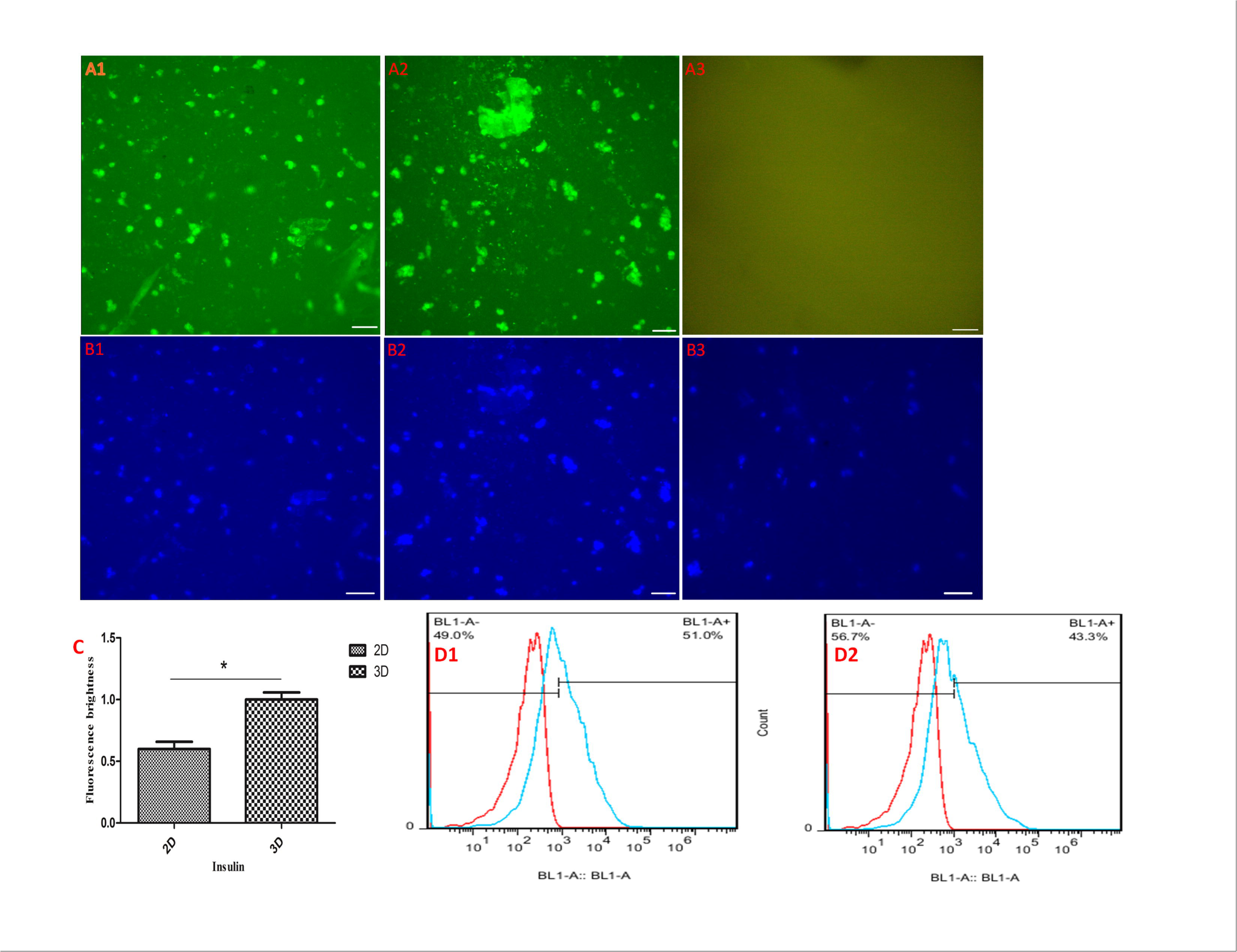
Immunocytochemical assays related to WJ-MSCs-derived IPCs. The cytoplasmic location of insulin in WJ-MSCs-derived IPCs at day 20 of the differentiation period in the 3D group (A2) and 2D group (A1), indicated by FITC (green). Nucleus staining was performed with DAPI (blue) in 2D (B1), 3D groups (B2) and along with secondary control test (B3). (A3) and (B3) were related to the secondary controls. Bar graph quantification of the immunocytochemistry assay (C). Each bar represents the relative value of insulin. Detection of insulin expression by flow cytometry in 3D (51.0%), (D1), and 2D (43.3%), (D2) groups. Scale bars are 100 μm.

### _Insulin and C-peptide release assay

At the end of the differentiation period, to evaluate the functional level of WJ-MSCs-derived IPCs, we exposed them to different concentrations of glucose solution. Next, the amount of insulin and C-peptide secreted by the WJ-MSCs-derived IPCs in the 3D and 2D experimental groups and the non-experimental control group were measured by the ELISA test. The measurement of insulin (Fig. 7A), and C-peptide (Fig. 7B) secretion from cultured cells in 2D and 3D experimental groups and the 2D control group was used to evaluate the functional level of the WJ-MSCs-derived IPCs at the end of the differentiation period. The secretion of insulin and C-peptide in response to different concentrations of glucose in the control group and also in response to the concentration of 5.5 mM glucose in the 2D and 3D experimental groups were not significant. The amount of insulin and C-peptide secretion in response to 15 and 25 mM glucose concentrations has increased significantly in the 3D experimental group compared with the 2D experimental group. The data showed that the PRP-PVP-PCL/PCL scaffold improves the maturity level and function of WJ-MSCs-derived IPCs.

**Fig. 7.**
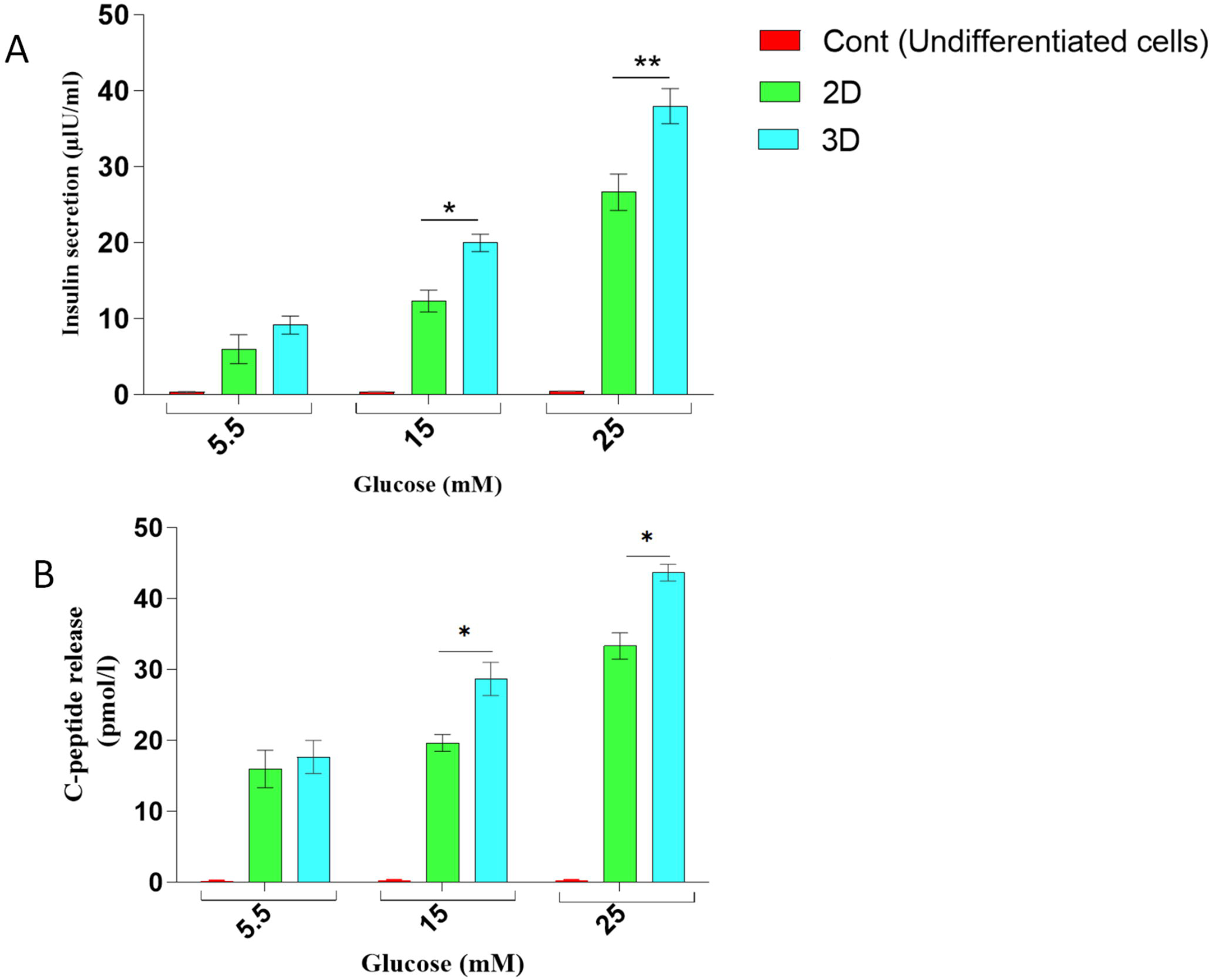
Assay of insulin and C-peptide release from WJ-MSCs-derived IPCs (A, B). Comparison of insulin and C-peptide secretion in response to different concentrations of glucose including 5.5, 15, and 25 mM in 2D, 3D, and control groups. Values are expressed as mean ± SEM. (n = 6). *p < .05 and **p < .01.

## Discussion

The ECM plays a vital role in the function and survival of differentiated cells. In 3D cell culture, due to mimicking the ECM, the survival and function of cells in 3D cell culture are significantly higher than in 2D culture(Aloysious and Nair, 2014; Mooranian et al., 2018). ENFs provide an ideal microenvironment similar to *in vivo* for cells because of their nano and porous structure. The strength and flexibility of the scaffold and the hydrophilicity of the scaffold surface are both necessary for cell attachment to the scaffold and cell migration and differentiation(Abazari et al., 2019; Maleki et al., 2021). PRP contains various types of growth factors that can help the differentiation of stem cells into IPCs. For example, b-FGF induces the differentiation of stem cells into pancreatic islets, EGF increases the expression of the Pdx1 gene, HGF increases the response to glucose by increasing insulin production in IPCs, and vascular endothelial growth factors A (VEGF-A) causes the formation of islet-shaped structures(Enderami et al., 2018a). We used PRP to improve the differentiation protocol. To fabricate the PRP-PVP-PCL/PCL (8%: 20%: 5%/ 15%) scaffold, we prepared the PRP-PVP solution. Then, the viscosity was increased with a 5% PCL solution. Electrospinning of 15% PCL solution with PRP-PVP-PCL solution in a layer-by-layer process helped in the phased release of PRP and increased the strength and stability of the scaffold. PVP caused the preservation of PRP and the hydrophilicity of the scaffold. In the study of Mohammad Foad Abazari et al., the burst of PRP release from polyvinyl alcohol (PVA)-chitosan-hydroxyapatite (HA)-PRP scaffold occurred after 24 hours(Abazari et al., 2018). In our study, it occurred after 48 hours; the scaffold lost 83% of its mass after 72 hours and burst release of PRP after 48 hours. Also, losing 83% of the scaffold mass was related to the PRP-PVP part of the scaffold. PRP, combined with PCL 5%, with the presence of PCL 15% as a reinforcement of the stability of the PRP-PVP-PCL/PCL scaffold, was gradually released. The contact angle test showed that PVP and PRP increased the hydrophilicity of the scaffold and compensated for PCL hydrophobicity. In the other study, to differentiate human induced pluripotent stem cells (hiPSs) into IPCs in 2D culture, PRP was added to the differentiation culture medium and MTT assay showed a significant decrease in the viability of hiPSs during the differentiation process. However, we showed an increase in cell survival on the scaffold containing PRP(Enderami et al., 2018a). In the study of Motaharesadat Hosseini et al., human adipose-derived Stem Cells (hADSCs)-derived IPCs in both 2D and 3D (Silk/polyethersulfone (PES) hybrid scaffold) experimental groups had morphology similar to pancreatic islet clusters as in our study(Hosseini et al., 2020). In the study of Samad Nadri et al., 21.7% of the cells in the 2D differentiation group were CJMSCs-derived IPCs, also in the concentration of 25 mM of glucose, the amount of insulin secretion in the 3D differentiation group was significantly higher than the 2D differentiation group. In our study, 43.3% of the cells in the 2D differentiation group were WJ-MSCs-derived IPCs and the amount of insulin secretion in 15 and 25 mM glucose concentrations was significantly increased in the 3D differentiation group(Nadri et al., 2018). In another study by Seyed Ehsan Enderami et al., the survival of hiPSCs on days 5 and 7 increased significantly in the 3D polyLLLlactic acid/polyvinyl alcohol (PLLA/PVA) group(Enderami et al., 2018a). This increase was not significant in our study. In this study, the expression of Pdx1, insulin, and GCG genes increased significantly in the 3D group, but the expression of the GLUT2 gene did not increase significantly. In our study, the expression of GLUT2 and other genes had a significant increase in the 3D differentiation group. In this study, there was a significant increase in the secretion of insulin and C-peptide at the concentration of 25 mM of glucose, but in our study at concentrations of 15 and 25 Mm of glucose showed a significant increase in the 3D differentiation group, this increase in the secretion of insulin and C-peptide in our study can be due to the significant increase in the expression of GLUT2 and a greater increase in the Pdx1 expression in the 3D differentiation group, which increased WJ-MSCs-derived IPCs maturity and function. In the study of Enderami et al., hADSCs were cultured on PVA scaffolds and the IPCs differentiation medium was optimized with PRP(Enderami et al., 2018b). Insulin and C-peptide secretion showed a significant increase in the 3D group only at the concentration of 25 mM glucose. In our study, observed 15 and 25 mM glucose. In our study PCL in the composition of the PRP-PVP-PCL/PCL scaffold, increased the level of IPCs maturity and function, because it provided a strong and flexible scaffold for the cells. The hydrophilicity of the scaffold was increased with PVP and PRP, and the gradual release of PRP from the scaffold was effective. In the study of Mohammad Ojaghi et al.,The expression of GLUT2 in hADSCs-derived IPCs did not show a significant increase in the 3D (PLLA/PVA) group, but this increase was significant in our study. The secretion of insulin and C-peptide in the 3D differentiation group at 15 and 25 mM glucose was the same as in our study, due to the lack of a significant increase in the expression of GLUT2, this significant increase in insulin and C-peptide secretion can be due to the increase in the number of hADSCs-derived IPCs due to the significant increase in the survival of hADSCs on the PLLA/PVA scaffold(Ojaghi et al., 2019). Cells must acquire their normal morphology to function normally. Researchers are trying to simulate and engineer the natural 3D conditions of cells to obtain optimally functional cells for therapeutic or research purposes. The ability of insulin-producing beta cells is considered in insulin production and regulation of insulin secretion in response to different concentrations of glucose. These cells obtain their normal morphology and optimal function in the structure of pancreatic clusters and in connection with other types of cells in pancreatic islets. The results of the analyses performed in our study showed that these characteristics were significantly higher in the 3D differentiation group than in the 2D differentiation group. Finally, the PRP-PVP-PCL/PCL scaffold as an ideal 3D matrix provides a suitable microenvironment for the engineering of pancreatic islets and promotes the maturity and functional level of WJ-MSCs-derived IPCs.

## Conclusion

Our study confirmed that the PRP-PVP-PCL/PCL scaffold as an innovative 3D matrix has provided an ideal 3D culture with the important features of strength, flexibility, and surface hydrophilicity. These features, together with the release of PRP, caused the proper WJ-MSCs attachment to the scaffold surface and the proliferation and differentiation of the stem cells. In our study, WJ-MSCs-derived IPCs expressed insulin as a specific marker of pancreatic beta cells in mRNA and protein levels, and they could regulate the amount of insulin secretion as the main function of insulin-producing cells in the glucose challenge test. All these results in the 3D differentiation group were significantly increased compared to the 2D differentiation group. In further studies, we can evaluate the quality of fiberized PRP after being released from the nanofibers scaffold. Also, the function of WJ-MSCs-derived IPCs with this method should be further investigated in vivo.

## Acknowledgments

This study was supported by research grants from the Mazandaran University of Medical Sciences (project number: 17642) and Sari Agricultural Sciences and Natural Resources University. We wish to thank Stem Cell Technology Research Center for the use of their laboratory facilities.

## References

1. Abazari, M.F., Nejati, F., Nasiri, N., Khazeni, Z.A.S., Nazari, B., Enderami, S.E. and Mohajerani, H.: Platelet-rich plasma incorporated electrospun PVA-chitosan-HA nanofibers accelerates osteogenic differentiation and bone reconstruction. Gene 720 (2019) 144096.

2. Abazari, M.F., Soleimanifar, F., Aleagha, M.N., Torabinejad, S., Nasiri, N., Khamisipour, G., Mahabadi, J.A., Mahboudi, H., Enderami, S.E. and Saburi, E.: PCL/PVA nanofibrous scaffold improve insulin-producing cells generation from human induced pluripotent stem cells. Gene 671 (2018) 50–57.

3. Aloysious, N. and Nair, P.D.: Enhanced survival and function of islet-like clusters differentiated from adipose stem cells on a three-dimensional natural polymeric scaffold: an in vitro study. Tissue Engineering Part A 20 (2014) 1508–1522.

4. Amable, P.R., Carias, R.B.V., Teixeira, M.V.T., da Cruz Pacheco, Í., Corrêa do Amaral, R.J.F., Granjeiro, J.M. and Borojevic, R.: Platelet-rich plasma preparation for regenerative medicine: optimization and quantification of cytokines and growth factors. Stem cell research & therapy 4 (2013) 1–13.

5. Bhattarai, D.P., Aguilar, L.E., Park, C.H. and Kim, C.S.: A review on properties of natural and synthetic based electrospun fibrous materials for bone tissue engineering. Membranes 8 (2018) 62.

6. Cakir, S., Gultekin, B.A., Karabagli, M., Yilmaz, T.E., Cakir, E., Guzel, E.E., Yalcin, S., Mortellaro, C. and Mijiritsky, E.: Histological evaluation of the effects of growth factors in a fibrin network on bone regeneration. Journal of Craniofacial Surgery 30 (2019) 1078–1084.

7. Choi, U.Y., Joshi, H.P., Payne, S., Kim, K.T., Kyung, J.W., Choi, H., Cooke, M.J., Kwon, S.Y., Roh, E.J. and Sohn, S.: An Injectable Hyaluronan–Methylcellulose (HAMC) Hydrogel combined with Wharton’s jelly-derived mesenchymal Stromal cells (WJ-MSCs) promotes degenerative disc repair. International journal of molecular sciences 21 (2020) 7391.

8. Enderami, S.E., Kehtari, M., Abazari, M.F., Ghoraeian, P., Nouri Aleagha, M., Soleimanifar, F., Soleimani, M., Mortazavi, Y., Nadri, S. and Mostafavi, H.: Generation of insulin-producing cells from human induced pluripotent stem cells on PLLA/PVA nanofiber scaffold. Artificial cells, nanomedicine, and biotechnology 46 (2018a) 1062–1069.

9. Enderami, S.E., Mortazavi, Y., Soleimani, M., Nadri, S., Biglari, A. and Mansour, R.N.: Generation of insulinLproducing cells from humanLinduced pluripotent stem cells using a stepwise differentiation protocol optimized with plateletLrich plasma. Journal of cellular physiology 232 (2017) 2878–2886.

10. Enderami, S.E., Soleimani, M., Mortazavi, Y., Nadri, S. and Salimi, A.: Generation of insulinLproducing cells from human adiposeLderived mesenchymal stem cells on PVA scaffold by optimized differentiation protocol. Journal of cellular physiology 233 (2018b) 4327–4337.

11. Haaf, F., Sanner, A. and Straub, F.: Polymers of N-vinylpyrrolidone: synthesis, characterization and uses. Polymer Journal 17 (1985) 143–152.

12. Himmler, M., Schubert, D.W., Dähne, L., Egri, G. and Fuchsluger, T.A.: Electrospun PCL Scaffolds as Drug Carrier for Corneal Wound Dressing Using Layer-by-Layer Coating of Hyaluronic Acid and Heparin. International Journal of Molecular Sciences 23 (2022) 2765.

13. Hosseini, M., DadashiLNoshahr, K., Islami, M., Saburi, E., Nikpoor, A.R., Mellati, A., MossahebiLMohammadi, M., Soleimanifar, F. and Enderami, S.E.: A novel silk/PES hybrid nanofibrous scaffold promotes the in vitro proliferation and differentiation of adiposeLderived mesenchymal stem cells into insulin producing cells. Polymers for Advanced Technologies 31 (2020) 1857–1864.

14. Jadhav, S., Nikam, D., Khot, V., Thorat, N., Phadatare, M.R., Ningthoujam, R., Salunkhe, A. and Pawar, S.: Studies on colloidal stability of PVP-coated LSMO nanoparticles for magnetic fluid hyperthermia. New Journal of Chemistry 37 (2013) 3121–3130.

15. Laidmäe, I., Aints, A. and Uibo, R.: Growth of MIN-6 Cells on Salmon Fibrinogen Scaffold Improves Insulin Secretion. Pharmaceutics 14 (2022) 941.

16. Mirzaei, A., Saburi, E., Enderami, S.E., Barati Bagherabad, M., Enderami, S.E., Chokami, M., Shapouri Moghadam, A., Salarinia, R., Ardeshirylajimi, A. and Mansouri, V.: Synergistic effects of polyaniline and pulsed electromagnetic field to stem cells osteogenic differentiation on polyvinylidene fluoride scaffold. Artificial cells, nanomedicine, and biotechnology 47 (2019) 3058–3066.

17. Liu, Y., Chen, X., Yu, D.-G., Liu, H., Liu, Y. and Liu, P.: Electrospun PVP-core/PHBV-shell fibers to eliminate tailing off for an improved sustained release of curcumin. Molecular Pharmaceutics 18 (2021) 4170–4178.

18. Lu, G., Li, S., Guo, Z., Farha, O.K., Hauser, B.G., Qi, X., Wang, Y., Wang, X., Han, S. and Liu, X.: Imparting functionality to a metal–organic framework material by controlled nanoparticle encapsulation. Nature chemistry 4 (2012) 310–316.

19. Maldonado, M., Huang, T., Yang, L., Xu, L. and Ma, L.: Human umbilical cord Wharton jelly cells promote extra-pancreatic insulin formation and repair of renal damage in STZ-induced diabetic mice. Cell Communication and Signaling 15 (2017) 1–13.

20. Maleki, H., Khoshnevisan, K., Sajjadi-Jazi, S.M., Baharifar, H., Doostan, M., Khoshnevisan, N. and Sharifi, F.: Nanofiber-based systems intended for diabetes. Journal of nanobiotechnology 19 (2021) 1–34.

21. Marx, R.E.: Platelet-rich plasma: evidence to support its use. Journal of oral and maxillofacial surgery 62 (2004) 489–496.

22. Mirtaghi, S.M., Hassannia, H., Mahdavi, M., HosseiniLkhah, Z., Mellati, A. and Enderami, S.E.: A novel hybrid polymer of PCL/fish gelatin nanofibrous scaffold improves proliferation and differentiation of Wharton’s jellyLderived mesenchymal cells into isletLlike cells. Artificial Organs 46 (2022) 1491–1503.

23. Mooranian, A., Negrulj, R., Takechi, R., Jamieson, E., Morahan, G. and Al-Salami, H.: Electrokinetic potential-stabilization by bile acid-microencapsulating formulation of pancreatic β-cells cultured in high ratio poly-L-ornithine-gel hydrogel colloidal dispersion: Applications in cell-biomaterials, tissue engineering and biotechnological applications. Artificial Cells, Nanomedicine, and Biotechnology 46 (2018) 1156–1162.

24. Musiał-Wysocka, A., Kot, M., Sułkowski, M., Badyra, B. and Majka, M.: Molecular and functional verification of Wharton’s jelly mesenchymal stem cells (WJ-MSCs) Pluripotency. International journal of molecular sciences 20 (2019) 1807.

25. Nadri, S., Barati, G., Mostafavi, H., Esmaeilzadeh, A. and Enderami, S.E.: Differentiation of conjunctiva mesenchymal stem cells into secreting islet beta cells on plasma treated electrospun nanofibrous scaffold. Artificial cells, nanomedicine, and biotechnology 46 (2018) 178–187.

26. Nekanti, U., Rao, V.B., Bahirvani, A.G., Jan, M., Totey, S. and Ta, M.: Long-term expansion and pluripotent marker array analysis of Wharton’s jelly-derived mesenchymal stem cells. Stem cells and development 19 (2010) 117–130.

27. Ojaghi, M., Soleimanifar, F., Kazemi, A., Ghollasi, M., Soleimani, M., Nasoohi, N. and Enderami, S.E.: Electrospun polyLlLlactic acid/polyvinyl alcohol nanofibers improved insulinLproducing cell differentiation potential of human adiposeLderived mesenchymal stem cells. Journal of Cellular Biochemistry 120 (2019) 9917–9926.

28. Pakfar, A., Irani, S. and Hanaee-Ahvaz, H.: Expressions of pathologic markers in PRP based chondrogenic differentiation of human adipose derived stem cells. Tissue and Cell 49 (2017) 122–130.

29. Perkins, B.A., Sherr, J.L. and Mathieu, C.: Type 1 diabetes glycemic management: Insulin therapy, glucose monitoring, and automation. Science 373 (2021) 522–527.

30. Pötter, N., Westbrock, F., Grad, S., Alini, M., Stoddart, M., Schmal, H., Kubosch, D., Salzmann, G. and Kubosch, E.: Evaluation of the influence of platelet-rich plasma (PRP), platelet lysate (PL) and mechanical loading on chondrogenesis in vitro. Scientific Reports 11 (2021) 1–11.

31. Abazari, M.F., Soleimanifar, F., Enderami, S.E., Nematzadeh, M., Nasiri, N., Nejati, F., Saburi, E., Khodashenas, S., Darbasizadeh, B. and Khani, M.M.: Incorporated bFGF polycaprolactone/polyvinylidene fluoride nanocomposite scaffold promotes human induced pluripotent stem cells osteogenic differentiation. Journal of cellular biochemistry 120 (2019) 16750–16759.

32. Riddell, M.C., Scott, S.N., Fournier, P.A., Colberg, S.R., Gallen, I.W., Moser, O., Stettler, C., Yardley, J.E., Zaharieva, D.P. and Adolfsson, P.: The competitive athlete with type 1 diabetes. Diabetologia 63 (2020) 1475–1490.

33. Rizal Syaidah, R., Aqsha, Z.M., Josephin, A. and Pakpahan, V.M.: Characterization, differentiation, and population doubling time of Wharton’s jelly mesenchymal stem cells (WJ-MSCs) in passage 5 and 8, AIP Conference Proceedings. AIP Publishing LLC, 2021, pp. 040002.

34. Saberzadeh-Ardestani, B., Karamzadeh, R., Basiri, M., Hajizadeh-Saffar, E., Farhadi, A., Shapiro, A.J., Tahamtani, Y. and Baharvand, H.: Type 1 diabetes mellitus: cellular and molecular pathophysiology at a glance. Cell Journal (Yakhteh) 20 (2018) 294.

35. Sánchez-Cardona, Y., Echeverri-Cuartas, C.E., López, M.E.L. and Moreno-Castellanos, N.: Chitosan/gelatin/pva scaffolds for beta pancreatic cell culture. Polymers 13 (2021) 2372.

36. Sun, B., Li, S., Zhang, H., Li, H., Zhao, C., Yuan, X. and Cui, Y.: Controlled release of Berberine Chloride by electrospun core/shell PVP/PLCL fibrous membranes. International Journal of Materials and Product Technology 37 (2010) 338–349.

37. Wang, L., Abhange, K.K., Wen, Y., Chen, Y., Xue, F., Wang, G., Tong, J., Zhu, C., He, X. and Wan, Y.: Preparation of engineered extracellular vesicles derived from human umbilical cord mesenchymal stem cells with ultrasonication for skin rejuvenation. ACS omega 4 (2019) 22638–22645.

38. Warshauer, J.T., Bluestone, J.A. and Anderson, M.S.: New frontiers in the treatment of type 1 diabetes. Cell metabolism 31 (2020) 46–61.

39. Yang, S.-J., Wang, X.-Q., Jia, Y.-H., Wang, R., Cao, K., Zhang, X., Zhong, J., Tan, D.-M. and Tan, Y.: Human umbilical cord mesenchymal stem cell transplantation restores hematopoiesis in acute radiation disease. American Journal of Translational Research 13 (2021) 8670.

40. Zhang, W., Ling, Q., Wang, B., Wang, K., Pang, J., Lu, J., Bi, Y. and Zhu, D.: Comparison of therapeutic effects of mesenchymal stem cells from umbilical cord and bone marrow in the treatment of type 1 diabetes. Stem Cell Research & Therapy 13 (2022) 1–14.

